# Learning to synchronize: Midfrontal theta dynamics during rule switching

**DOI:** 10.1101/2020.06.01.127175

**Authors:** Pieter Verbeke, Kate Ergo, Esther De Loof, Tom Verguts

## Abstract

In recent years, several hierarchical extensions of well-known learning algorithms have been proposed. For example, when stimulus-action mappings vary across time or context, the brain may learn two or more stimulus-action mappings in separate modules, and additionally (at a hierarchically higher level) learn to appropriately switch between those modules. However, how the brain mechanistically coordinates neural communication to implement such hierarchical learning, remains unknown. Therefore, the current study tests a recent computational model that proposed how midfrontal theta oscillations implement such hierarchical learning via the principle of binding by synchrony (Sync model). More specifically, the Sync model employs bursts at theta frequency to flexibly bind appropriate task modules by synchrony. 64-channel EEG signal was recorded while 27 human subjects (Female: 21, Male: 6) performed a probabilistic reversal learning task. In line with the Sync model, post-feedback theta power showed a linear relationship with negative prediction errors, but not with positive prediction errors. This relationship was especially pronounced for subjects with better behavioral fit (measured via AIC) of the Sync model. Also consistent with Sync model simulations, theta phase-coupling between midfrontal electrodes and temporo-parietal electrodes was stronger after negative feedback. Our data suggest that the brain uses theta power and synchronization for flexibly switching between task rule modules, as is useful for example when multiple stimulus-action mappings must be retained and used.

**Significance Statement:** Everyday life requires flexibility in switching between several rules. A key question in understanding this ability is how the brain mechanistically coordinates such switches. The current study tests a recent computational framework (Sync model) that proposed how midfrontal theta oscillations coordinate activity in hierarchically lower task-related areas. In line with predictions of this Sync model, midfrontal theta power was stronger when rule switches were most likely (strong negative prediction error), especially in subjects who obtained a better model fit. Additionally, also theta phase connectivity between midfrontal and task-related areas was increased after negative feedback. Thus, the data provided support for the hypothesis that the brain uses theta power and synchronization for flexibly switching between rules.

---

Switching between rules is key to function in a complex and rapidly changing environment. For instance, when at the pub with friends, our behavior is likely guided by different social rules than at work. However, when the boss suddenly walks into the pub, this requires to flexibly switch between these two sets of social rules. Importantly, an empirically valid model that explains how the human brain mechanistically deals with such switches, remains lacking.

In experimental settings, this cognitive flexibility in rule switching is typically tested in a reversal learning setup (Izquierdo, Brigman, Radke, Rudebeck, & Holmes, 2017). Here, agents must learn task rules, each consisting of a collection of stimulus-action mappings. During the task, these rules are regularly reversed. One popular framework to explain performance during reversal learning tasks is the Rescorla-Wagner model (RW; Rescorla & Wagner, 1972; Widrow & Hoff, 1960). Here, on every trial, obtained reward is used to update the value of active stimulus-action mappings. By learning fast, the agent can flexibly deal with changes in task rules. However, when feedback is probabilistic (e.g., Cools, Clark, Owen, & Robbins, 2002), this approach experiences difficulties. Specifically, a high learning rate will lead agents to “chase the noise” introduced by probabilistic feedback. In contrast, a low learning rate increases robustness against noise, but decreases flexibility on rule switches. Thus, some researchers have proposed that learning rate should be adaptive (e.g., Bai, Katahira, & Ohira, 2014; Behrens, Woolrich, Walton, & Rushworth, 2007; Silvetti, Vassena, Abrahamse, & Verguts, 2018). In this adaptive learning rate (ALR) proposal, agents track rule switches by comparing an estimate of reward probability to received reward. Consistently high prediction errors indicate that the underlying rule has changed, and learning rate should be increased. More fundamentally however, irrespective of learning rate flexibility, both RW and ALR frameworks assume that, on every rule reversal, old information is overwritten. Especially for more complex problems, this is inefficient, as is demonstrated by the problem of catastrophic forgetting in artificial neural networks (French, 1999).

To overcome catastrophic forgetting, separate task rules may be stored (Saez, Rigotti, Ostojic, Fusi, & Salzman, 2015; Wilson, Takahashi, Schoenbaum, & Niv, 2014). This poses a new problem of keeping track which task rule is currently relevant. Recent fMRI research focusing on this hierarchical approach toward reversal learning has pointed to midfrontal cortex as the responsible neural structure for keeping track of the current task rule (Wilson et al., 2014). However, how midfrontal cortex mechanistically coordinates neural communication in switching between task rules, remains an open question.

This question was recently addressed by a novel computational framework of hierarchical learning (Verbeke & Verguts, 2019). This Sync model retains separate mappings for every task rule, and keeps track of rule reversals by calculating prediction error (e.g., Holroyd & McClure, 2015), thus avoiding catastrophic forgetting. In order to guide neural communication between areas holding the appropriate mappings, the model relies on binding by synchrony (BBS; Fries, 2005, 2015; Gray & Singer, 1989; Womelsdorf et al., 2007) in theta frequency (4-8 Hz). Specifically, midfrontal theta oscillations synchronize neuronal activity along task-relevant pathways. Thus, task-relevant neurons can communicate and learn, while stability is achieved in currently irrelevant pathways.

The current study empirically tests this Sync model (Fig 1A). For this purpose, the model is fitted on data of subjects performing a probabilistic reversal learning paradigm, and empirically compared to alternative models (Bai et al., 2014; Rescorla & Wagner, 1972). Then, Sync model simulations provided several predictions for EEG measured during the task, specifically in theta frequency (model-driven EEG predictions). First, a linear relationship between midfrontal theta power and negative prediction errors was predicted, especially in subjects with good behavioral Sync model fit. Second, a peak of midfrontal theta power was predicted for data locked to rule switches. Third, phase-coupling between midfrontal and posterior electrodes was predicted to be stronger after negative feedback.

**Fig 1.**
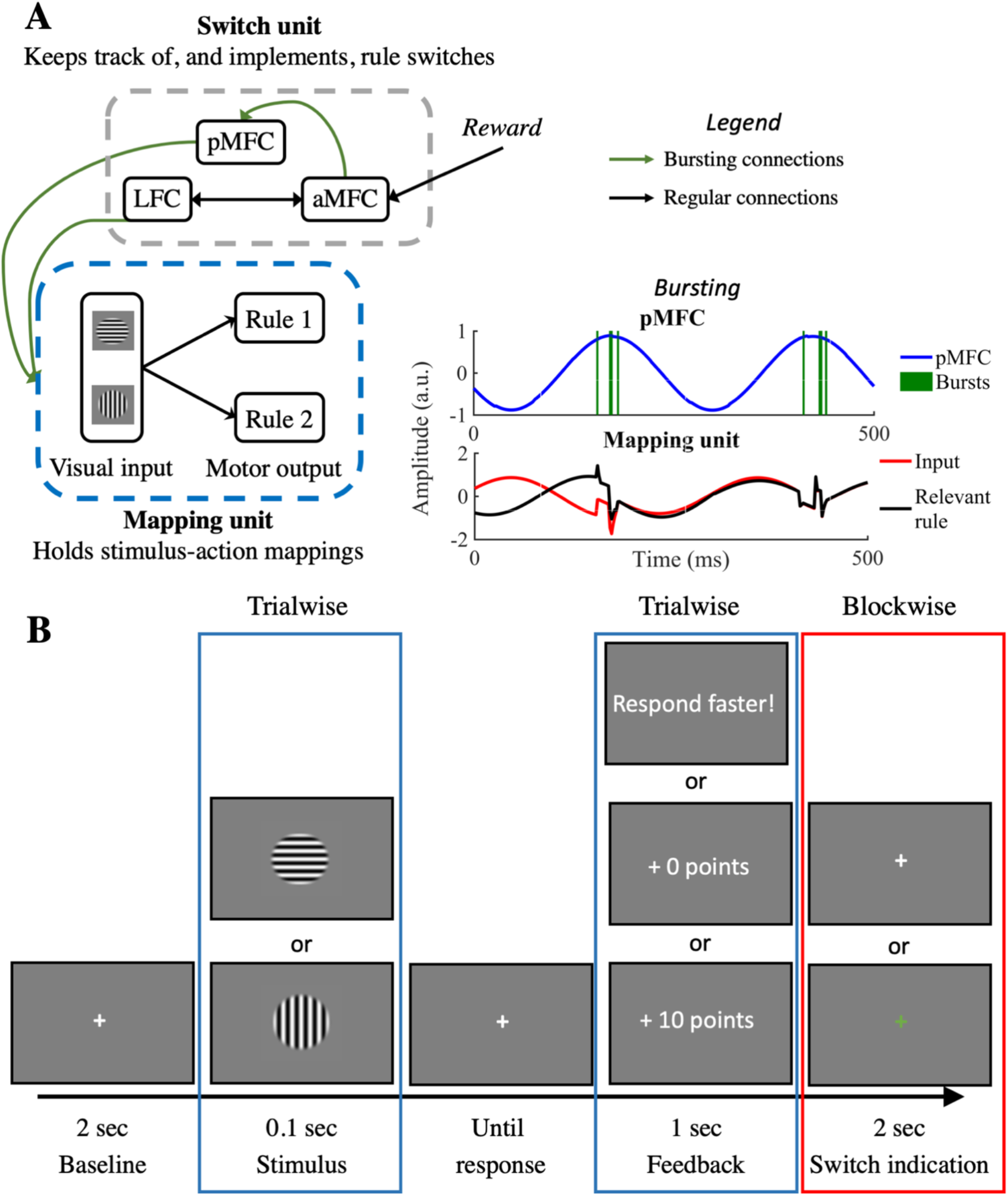
Methods. *A: The Sync model*. The left diagram represents a schematic overview of the Sync model. In the lower right corner, a detailed illustration shows how bursts originating in the pMFC synchronize task-relevant areas in the Mapping unit (see Verbeke & Verguts, 2019 for detailed explanation). *B: The task*. The time course of one trial in the experimental paradigm is shown. Elements highlighted by a blue rectangle, such as the presented stimulus and feedback, are manipulated on a trial-by-trial basis. Elements highlighted by the red rectangle are manipulated blockwise. Here, the fixation cross after feedback was green in one experimental block (half of all trials). In this reporting block, subjects had to press the space bar during this period if they thought the rule had switched.

## Materials and Methods

### Materials

The experiment was run on a Dell Optiplex 9010 mini-tower running PsychoPy software (Peirce et al., 2019). Electrophysiological data were recorded using a BioSemi ActiveTwo system (BioSemi, Amsterdam, Netherlands) with 64 Ag/AgCl electrodes arranged in the standard international 10–20 electrode mapping (Jasper, 1958), with a posterior CMS-DRL electrode pair. Two reference electrodes were positioned at the left and right mastoids. Eye movements were registered with a pair of electrodes above and below the left eye and two additional electrodes at the outer canthi of both eyes. EEG signals were recorded at a sampling rate of 1024 Hz.

Models were fitted using the differential evolution method of the SciPy (version 1.4.1) package in Python (version 3.7.6). Other behavioral analyses were done using R software (R Core Team, 2017). The electrophysiological data were preprocessed in MATLAB R2016b (The MathWorks Inc., 2016) using an EEGLAB preprocessing pipeline (Delorme & Makeig, 2004). Also for simulations of the Sync model MATLAB R2016b was used.

### Code and Data Accessibility

All code used to provide the results described in the current paper is provided at https://github.com/CogComNeuroSci/PieterV_public/tree/master/Reversal_learning. At publication, also the data will be made freely accessible at https://osf.io/wt36f/.

### Experimental Task

Both the model (27 simulations) and human subjects (*N* = 27) performed a probabilistic reversal learning task (see Fig 1B). Agents had to learn task rules consisting of two stimulus-action mappings which were regularly reversed during the task. Every trial started with a centrally presented white fixation cross for 2000 milliseconds. Then the stimulus was presented for a period of 100 milliseconds. This stimulus was a centrally presented circular grating with a raised-cosine mask and a size of 7 visual degrees. The grating was either vertically or horizontally oriented. After stimulus presentation, the screen turned blank until response. Responses were given by pressing the ‘f’- (left) or ‘j’ (right) key on an azerty keyboard. In task rule 1, the horizontal stimulus mapped to a left response and the vertical stimulus to the right response; this was reversed for task rule 2. During the task (480 trials), 15 rule switches were introduced. These rule switches occurred at random (uniform distribution from 15 to 45 trials after the previous task switch). After response, probabilistic feedback was presented in the center of the screen. This feedback consisted of ‘+10 points’ for rewarded trials, ‘+0 points’ for unrewarded trials or ‘Respond faster!’ when response times (RT) were slower than 1000 milliseconds. Subjects had an 80% probability of receiving reward feedback after correct responses and 20% after incorrect responses. After feedback, the fixation cross appeared again for another 2000 milliseconds. Crucially, the experiment was divided into two experimental blocks (240 trials each). In one block, the reporting block, the post-feedback fixation cross was presented in green. During this period, subjects were instructed to press the space bar if they thought the task rule had switched. The purpose of this approach was to obtain an indication of when the subject reached his or her own ‘Switch threshold’, as happens in the Sync model. This was only done during one block, so critical changes due to this difference in task structure could be checked. The order of the two blocks was counterbalanced across subjects. In between blocks, as well as three times within a block, subjects were allowed a short break. This break could only occur if there was no rule switch within 10 trials from the break.

### Human Testing Procedure

34 subjects participated in this study, 7 subjects were removed because of either technical problems with the EEG recording (4) or an inability to give a correct response on more than 2/3 of the trials (3), resulting in *N* = 27 (N_male_ = 6, N_female_ = 21). Subjects were told they would receive €25 for their participation, with a possibility to earn up to €3 extra reward depending on their performance.

Before starting the task, the subject had to go through two short practice sessions with gratings that were tilted 45° to the left or to the right relative to a vertical line. In the first practice session, the subject performed 30 trials with only one task rule. Here, the goal was to let the subject get acquainted with the general paradigm and learn a task rule through probabilistic feedback. Subjects were only allowed to continue to the second practice session if they performed above chance level (50%) and could report the correct task rule to the experimenter. If not, they performed this practice session again. In the second practice session, subjects performed 60 trials of the task with 3 rule switches and with the post-feedback green fixation cross (as in the reporting block). In this session, subjects pressed the space bar to indicate a task switch and received feedback for each press. The press was considered correct if subjects responded within 10 trials from the actual rule switch. They were allowed to continue to the next task if they were able to perform above chance level and had at least 1 correct indication of a rule switch. After successfully performing both practice sessions, subjects performed 480 trials of the actual task.

### Behavioral Analyses

To check for differences between the reporting block (green fixation cross) and the non-reporting block (see Experimental Task and Fig 1B), paired t-tests were performed for both accuracy and RT, depending on experimental block. In order to deal with the skewed distribution of RT, the natural log of RT was used for all analyses. Additionally, trials with too late responses (RT > 1000 milliseconds; 2.11% of all data) were excluded for both behavioral and EEG analyses.

### Model Analyses

More extensive analyses of behavioral data were done with a model-based approach. Current work aims to test the Sync model (Verbeke & Verguts, 2019), but two baseline models were fitted as well. In the following section, we first provide a detailed overview of the Sync model, followed by a description of all three models that were fitted on behavioral data. Then, we describe how model fit was evaluated.

#### The Sync model

An overview of model architecture is provided in Fig 1A. The Sync model consists of two units, the Mapping and Switch unit. The Mapping unit contains a classic network with 2 layers (visual input and motor output). Here, weights are adapted with the RW algorithm (Widrow & Hoff, 1960). In the Sync model, 4 nodes (2 for each response option) at the motor output layer, are divided in 2 rule modules, one for each task rule. Hence, as in (Wilson et al., 2014), the Mapping unit holds separate stimulus-action mappings for each task rule. In addition, a Switch unit forms a hierarchically higher network modeled after primate prefrontal cortex. This Switch unit keeps track of switches in task rule. Specifically, the Switch unit consists of the lateral frontal cortex (LFC), posterior medial frontal cortex (pMFC) and anterior midfrontal cortex (aMFC). Here, the LFC holds pointers (e.g., Botvinick et al., 2001; Cohen, Dunbar, & McClelland, 1990) that indicate which rule should be synchronized in the Mapping unit. Since BBS implements gating, allowing efficient communication between synchronized nodes and blocking communication between non-synchronized nodes (Fries, 2005, 2015), the agents’ behavior will be guided by the synchronized rule. This synchronization process is then executed by the binding by random bursts principle (Springer & Paulsson, 2006; Verguts, 2017; Zhou, Chen, & Aihara, 2005). In the Sync model, a theta-frequency-paced signal produced in the pMFC is responsible for sending these bursts (see Verbeke & Verguts, 2019; Verguts, 2017 for details). The aMFC contains a neural network (for simplicity not shown in Fig 1A) that is adapted from previous work (Silvetti, Seurinck, & Verguts, 2011). Here, again RW learning is employed but on a hierarchically higher level. More specifically, the aMFC learns an expected reward (*V*) for the currently used rule module (see Equation (6)). This expected reward is compared to an external reward signal (*Rew*; Reward in Fig 1A) in order to compute prediction errors. The negative prediction error signal is propagated to both the Accumulator neuron (within the aMFC neural network) and to pMFC. A single negative prediction error increases (via bursting) the power of the theta signal in pMFC (bursting connection in Fig 1A; see Verbeke & Verguts, 2019 for details). Instead, the Accumulator neuron evaluates the prediction error signal on a slower time scale (see also Holroyd & McClure, 2015), and thus requires multiple prediction errors before activation in the Accumulator neuron reaches its Switch threshold (see Equation (5)). When this happens, aMFC signals the need for a switch to the LFC. Correspondingly, the LFC will change the signal to the Mapping unit, and synchronize another rule module. In sum, bursts received by the Mapping unit are the result of a cooperation between LFC and pMFC. The pMFC determines the intensity of theta bursts, while the LFC determines which task rule in the Mapping unit is susceptible to the bursts. For further details see (Verbeke & Verguts, 2019).

All nodes in the visual input and motor output layer of the Mapping unit as well as the pMFC are oscillatory nodes. In line with previous work (Verguts, 2017), oscillatory nodes consist of neuronal triplets. The neural triplet contains one excitatory-inhibitory pair of phase code neurons (E, I) and a rate code neuron. Here, excitatory neurons are updated by

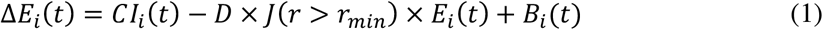

where Δ*E*(*t*) = *E*(t + Δ*t*)–*E*(*t*); and inhibitory neurons are updated by

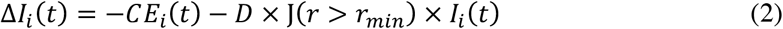

Here, phase code neurons will oscillate at a frequency of *C*/2*π*. In the pMFC, which executes top-down control by sending bursts, activity oscillates at theta (6 Hz) frequency, in line with suggestions of previous empirical work (Cavanagh & Frank, 2014; Womelsdorf, Johnston, Vinck, & Everling, 2010). Different from our previous modelling work, theta frequency was used in the Mapping unit (see Discussion) as well. Because bursts (*B*(t)) lead to a significant increase of power, a radius parameter (*r*_*min*_) is implemented in order to attract power (*r*) back to baseline after a burst. Since continuously high pMFC power is computationally suboptimal and empirically implausible (Holroyd, 2016), power in the pMFC was attracted towards a smaller radius, *r*_min_ = .50, than in the Mapping unit, r_min_ = 1. How fast oscillations decay to baseline is determined by a damping parameter (*D*) which was set to *D* = .30 in the Mapping unit. Since the pMFC not only receives bursts but also sends them, a slower decay *D* = .01 was implemented here to allow a sufficient activity window (∼ 500 ms/3 theta cycles) for bursts to be sent. In order to reduce model complexity, no oscillations were used in the LFC and aMFC. For a full description of model dynamics see Verbeke & Verguts, (2019).

Thus, in the Sync model, on every trial multiple time steps were simulated in which oscillations occurred. Here, motor nodes accumulate activation over time. The motor node with the maximal accumulated activation over time, was considered as the model response. Values of stimulus action pairs (*Q*) in each rule module (*R*) are updated by

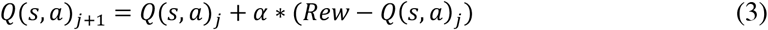

in which *α* is the Mapping learning rate and *Rew* is the reward received by the agent.

As described above, the Sync model has an additional Switch unit which adds a hierarchical learning algorithm on top of the RW (fixed learning rate) algorithm in the Mapping unit. This Switch unit evaluates whether there was a rule switch. More specifically, it learns a value (*V*) for every rule module (*R*) by

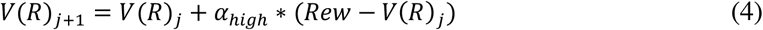

in which *α*_*high*_ is the hierarchically higher Switch learning rate. The difference between the expected value *V*(*R*) in Equation (6) and the obtained *Rew* (i.e., the prediction error) is accumulated in the Accumulator neuron (*A*) via

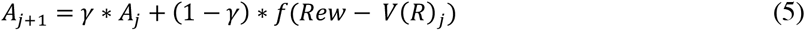

Since switches are only required when negative feedback occurs, the Accumulator neuron was selective for negative prediction errors. Specifically, *f*(*Rew* − *V*(*R*)) = −(*Rew* − *V*(*R*)) when the prediction error is negative and *f*(*Rew* − *V*(*R*)) = 0 when the prediction error is positive. Here, *γ* is the Cumulation parameter which determines how strongly the Accumulation neuron is affected by a single prediction error. While a low Cumulation parameter causes the agent to strongly weigh single prediction error and therefore regularly switch between rule modules, a high Cumulation parameter implements a more conservative approach. When the Accumulator neuron reaches a Switch threshold of .5, the model will switch to another rule module (*R*) in the Mapping unit.

#### Behavioral data fitting

For behavioral data fitting only, the full Sync model was simplified by introducing a hard gating process between task rules instead of BBS and a softmax response selection mechanism described by

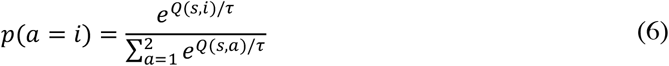

in which *Q(s,a*) is the value of a given stimulus-action pair (*s, a*) and *τ* is the temperature parameter which determines how strongly the agent explores different actions (*i*). This allowed to skip the loop of 1500 timesteps every trial, which was needed to simulate oscillations (see Equations (1) and (2)). We refer to this model as the behavioral Sync (bSync) model.

On top of the bSync model, two other models were fitted as well. The RW and ALR model are both restricted to only the Mapping unit (with one rule module). Both models use a response selection mechanism as described by Equation (6) and learn stimulus-action pairs by Equation (3). Importantly, the RW model (Rescorla & Wagner, 1972) had a constant learning rate while the ALR model (Bai et al., 2014), was implemented with an adaptable learning rate. Here, the Mapping learning rate is updated on every trial by

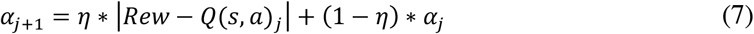

in which *η* determines how strongly the learning rate is influenced by the current difference between *Rew* and *Q* (lower-level prediction error).

#### Model evaluation

For each subject, the goodness of fit of these three models on the behavioral data was compared by using three measures. The log-likelihood (*LL*)

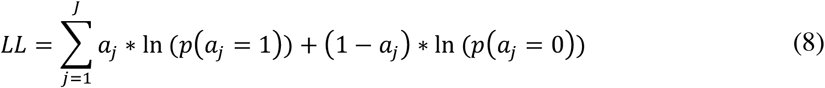

in which *p*(*a*) is the probability of the given action (see Equation (6)) and *J* represents the number of trials. The Akaike information criterion (AIC) uses this *LL* but includes a penalty for the number of parameters (*k; k* = 2 for RW, *k* = 3 for ALR and *k* = 4 for bSync) that were used in the model:

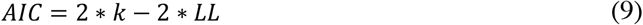

From this AIC, AIC weights (*wAIC*) can be derived which allows to make a relative comparison between the model fit of the three different models. These wAIC values are computed as

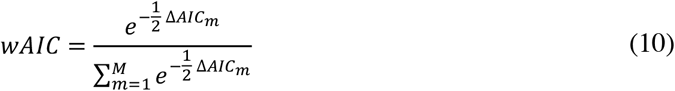

in which *M* is the number of models that are compared (*M* = 3) and

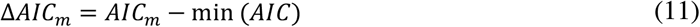

Here, min(AIC) is the lowest AIC value out of the three models for that subject. Thus, Equation (10) results in a wAIC value for each model. The sum of all three wAIC values is 1 and models with higher wAIC values provide a better fit to the data.

#### Simulations

In order to provide hypotheses for EEG data, 27 simulations of the full Sync model were performed. For all simulations, the same parameter values were used. These parameter values were sampled from the distribution of best fitting parameter values of the bSync model so that overall accuracy of model simulations (*M* = 78.00%, *SD* = 1.30) closely resembled accuracy of subjects (*M* = 76.80%, *SD* = 4.91). This resulted in a Mapping learning rate (*α*) of .8, a Switch learning rate (*α*_*high*_) of .1 and a Cumulation parameter (*γ*) of .3. The full Sync model did not use a Temperature (*τ*) parameter; instead, the synchronization procedure introduces noise, which also introduces some randomness in behavior. The Switch threshold was always fixed to .5. Trials were simulated as a fixed period of 500 ms in which the visual layer received stimulation. After this period, the response node with the highest maximum activation was registered as the response of the model. Thereafter, 1500 ms of inter-trial interval was simulated in order to provide a post-feedback period that could be analyzed in the same way as the empirical data. All other aspects of the task, such as the frequency and timing of rule switches, were the same for the model as for the human subjects.

#### Power Analyses

Time-frequency decomposition was performed on the excitatory neuron (see Equation (1)) within the neuronal triplet of the model’s pMFC node in the model. Complex Morlet wavelets were used for frequencies between 2 and 48 Hz defined in 25 logarithmically spaced steps. For each frequency, between 3 and 8 cycles were used, also defined in 25 logarithmically spaced steps. Power was extracted as the squared absolute value of the time-frequency decomposed signal. In order to locate activity that was specific to feedback processing, the difference between power in trials with negative feedback and trials with positive feedback was computed. For simplicity, we selected the 2.5% most positive values as a cluster of interest. This cluster contained one group of data points in theta frequency and approximately 250-500 ms after feedback (see Verbeke & Verguts, 2019 for timing details). On every trial, the mean power in this cluster was computed and entered in the consecutive analyses. Since a negative prediction error in the model increases activity of the pMFC, we performed a linear regression of cluster power with prediction error as the independent variable. To test our first hypothesis, that this relationship was specific to negative prediction error, a second regression model was used that also included the interaction between prediction error and reward. The second hypothesis states that because negative prediction errors are strongest at the moment of a rule switch, a peak of post-feedback theta power should be found when data is locked to rule switches. To investigate this, we extracted power from the model cluster in trials within a 31-trial window around the rule switch (−15 to +15). The time course (one data point for each of the 31 trials) that resulted after averaging over all (15) rule switches and all (27) simulations was then used as a regressor in a linear regression with data from the empirical clusters.

#### Phase Analyses

Our third hypothesis stated that phase-coupling between pMFC and model nodes in the Mapping unit was stronger after negative feedback. Specifically, theta power in the model pMFC increases after negative prediction errors. When there is sufficient power in the pMFC, it will increase synchronization in the Mapping unit (posterior/lateral task-related regions, e.g., pre-motor or visual areas). For this purpose, the pMFC uses binding by random bursts (Verguts, 2017). Here, the pMFC will send bursts to the Mapping unit at specific phases. Thereby it will shift the phase of neurons in the Mapping unit (see Verbeke & Verguts, 2019 for details). This leads to phase shifts in these lower pre-motor or visual task-related areas, and a short period of phase-alignment between these task-related areas in the Mapping unit and pMFC. Phase was extracted in all model nodes by taking the angle of the Hilbert transform of the raw signal. For simplicity the model was implemented without inter-areal delays. Furthermore, in contrast to analyses on the empirical EEG data (see Equation (12)), control for volume conduction was not needed, so the regular phase locking value (PLV; Lachaux, Rodriguez, Martinerie, & Varela, 1999) was computed between the model pMFC and the nodes in the motor layer of the Mapping unit. This PLV was then averaged over all 4 motor nodes and the time period included in the power cluster (∼250-500 ms post feedback).

### EEG Analyses

#### Preprocessing

The data were re-referenced offline to the average of the mastoid electrodes. Breaks or other offline periods were manually removed. Particularly noisy electrodes were interpolated between neighboring electrodes on all timesteps. For three subjects, one electrode was interpolated; for another three subjects we had to interpolate two electrodes; because of a bridge, one subject needed interpolation for five posterior electrodes. Additionally, activity was band-pass filtered between 1 and 48 Hz in order to remove slow drifts and line noise of 50 Hz. Eyeblinks and other motor-related noise components were removed through EEGLAB independent component analysis (ICA). After ICA-removal, the data was epoched, once locked to feedback onset, and once to stimulus onset. The epochs based on stimulus onset were used to extract baseline activation, which was −1500 to −500 ms relative to stimulus onset. This baseline activity was subtracted from all epochs. After epoching, on average 7.5% of epochs were removed by applying an amplitude threshold of −500 to 500 mV and an improbability test with 6 standard deviations for single electrodes and 2 standard deviations for all electrodes, as described in Makoto’s preprocessing pipeline (Makoto, 2018). Before time-frequency analyses, data was also downsampled to 512 Hz.

#### Time-frequency Decomposition

Time-frequency decomposition was based on code from (Cohen, 2014). Similar to model analyses, complex Morlet wavelets were used for frequencies between 2 and 48 Hz defined in 25 logarithmically spaced steps. For each frequency, between 3 and 8 cycles were used, also defined in 25 logarithmically spaced steps.

#### Power Computation

A baseline correction was applied by dividing the power estimates for each subject, electrode and frequency by the average baseline activity (−1500 ms to −500 ms from stimulus onset) across all 480 trials. Finally, the baseline-corrected data underwent a decibel conversion. Before final analyses, also trials with late responses were removed from the data.

#### Power Cluster Analyses

Similar to model analyses, we were interested in activity selective for feedback. Hence, a contrast between Z-scored power in trials with negative feedback and trials with positive feedback was computed. On these values, a non-parametric clustering procedure was applied (Maris & Oostenveld, 2007). The distribution of statistics was computed. On each side of the distribution (two-sided test), the 1% most extreme values were entered into the clustering analysis. From these, we clustered adjacent neighbors in the channel, frequency and time domains. To calculate our cluster-level statistic, we multiplied the number of items (i.e., (channel, frequency, time) points) in the cluster with the largest statistic of that cluster (see also Maris & Oostenveld, 2007). A significance threshold of 5% was imposed on the subsequent non-parametric permutation test with 1000 iterations. Clusters that survived this permutation test were taken into the consecutive analyses. As an exploratory analysis, we aimed to link individual differences in behavioral model fit to EEG data; for that purpose, we extracted the mean cluster statistic for each subject, and ran a Spearman rank correlation of these statistics with wAIC of the bSync model obtained in the behavioral model fitting procedure.

#### Midfrontal Theta Power and Prediction Error

The Sync model uniquely yields specific EEG predictions, to which we now turn. To test the first model-driven EEG hypothesis of a relation between theta power and prediction errors, we first extracted a measure of prediction error for every subject on every trial by simulating the bSync model. Importantly, this prediction error was extracted from the learning process on the hierarchically higher level in the Switch unit (Equation (4)), not the lower-level learning process in the Mapping unit (Equation (3)). This measure of prediction error was then used in a trial-by-trial linear mixed effects model as a predictor for the Z-scored power of every cluster (averaged across all (time, electrode, frequency points) in the cluster), that survived the feedback-locked analysis described above. Here, a random intercept for every subject was included and a fixed slope (i.e., the prediction error). Because the Sync model predicted different relationships for positive prediction errors and negative prediction errors, also the interaction between prediction errors and reward was tested. Additionally, in order to explore whether the individual differences in wAIC influenced the interaction between prediction errors and reward, also a three-way interaction between prediction error, reward and wAIC was tested. For these purposes, three regression models were fitted: One in which only prediction error was included as regressor, one in which both prediction error and the interaction between prediction error and reward were included as regressors, and finally a third model in which the main effect, the two-way interaction, and an extra three-way interaction between prediction error, reward and wAIC were included as regressors. These regression models were then compared via ANOVA.

#### Rule Switch Locking

A second model-driven EEG hypothesis considers theta power locked to the moment of a rule switch. For this analysis, EEG data of 31 trials around the rule switch (−15 to +15 trials, including the rule switch trial itself) were extracted. On these trials, the mean power within each cluster selective for feedback was computed. This data was then again averaged over all trials at a specific distance (−15 to +15) from switch, giving us a time course of mean cluster-power from −15 trials before rule switch to 15 trials after rule switch for every subject. On each time point, a 99.84% confidence interval (CI) was computed based on a Bonferroni correction for multiple comparisons (100-(5/31)). This confidence interval was compared to a baseline power. Baseline power was computed based on the mean power in this cluster, averaged over all trials that were more than 15 trials removed from the rule switch.

As the rule switch trial, we considered in separate analyses both the actual rule switch and the subjective indication of a rule switch. Hence, power close to a rule switch was compared with the mean power of trials that were far from the rule switch. When the confidence interval did not include the baseline value, power on this trial was considered as significantly deviating from baseline. Additionally, we aimed to investigate the similarity between the data pattern predicted by the model and the empirical data. For this purpose, data from the bSync model simulations (see above for details) was used as a linear regressor for the empirical data. Also for this hypothesis, an extra analysis was performed to investigate whether wAIC had an influence on the observed effect. Here, we extracted subject data on trials of which cluster power significantly deviated from baseline and used this data as a dependent variable in a linear regression with wAIC.

#### Midfrontal-Posterior Phase-Coupling Analyses

For the third model-driven EEG hypothesis, we considered all midline electrodes (10) as seed and other electrodes (54) as receiver in the phase connectivity analyses. Because we were interested in phase-locking related to rule modules conveying the correct response, all data was lateralized with respect to the correct response. All data ipsi-lateral to the correct response was brought to the left electrodes; all contra-lateral data was brought to the right electrodes. The iPLV (Bruña, Maestú, & Pereda, 2018) was computed between all midline electrodes and all lateral electrodes for every time point in the feedback-locked data. This iPLV measure was computed by the following equation

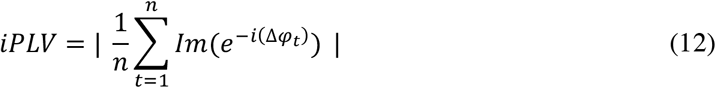

which computes the average phase angle (φ) difference over trials (t). By only looking at the imaginary (Im) part of this phase angle difference, phase differences of zero are eliminated. Hence, volume conduction effects are excluded, because such volume conduction effects are represented in zero-phase differences (Bruña et al., 2018; Nolte et al., 2004). Again, a non-parametric cluster algorithm was performed on the contrast between iPLV for trials with negative versus positive feedback (note that the fact that our effect of interest compares negative versus positive feedback, also safeguards against possible volume conduction effects). For this analysis, only data of one midline electrode was used. More specifically, we checked on which of the 10 midline electrodes the mean contrast in the theta frequency (4-8 Hz) reached a maximum. This was in the FCz electrode, hence only iPLV between FCz and all lateral electrodes were entered in the clustering algorithm. As for power, an exploratory analysis was performed in which we extracted the mean cluster statistic for each subject, and ran a Spearman rank correlation of these statistics with wAIC of the bSync model obtained in the model fitting procedure.

## Results

### Behavioral Data

Overall, participants had a mean accuracy of 76.80% (*SD* = 4.92%) and a mean RT of 544ms (*SD* = 71.31 ms). A paired t-test confirmed that there were no significant differences between the experiment block in which subjects had to indicate when a task switch happened or when they did not have to indicate this (see Materials and Methods for details), neither in accuracy (*t*(26) = .029, *p* = .977), nor in RT (*t*(26) = −1.290, *p* = .208).

#### Model Analyses

The distribution of all fitted parameter values for each model is given in Fig 2A. Goodness of fit measures are summarized in Table 1. Here, log-likelihood was highest (best) for the bSync model, lowest for the ALR model, with the RW model in between. When a penalty for model complexity was applied (AIC, wAIC), the RW and bSync models performed approximately equal. Importantly, wAIC results indicated significant differences across individuals. As illustrated in Fig 2B, subjects could be roughly divided into three groups based on the wAIC. In one group (8 subjects), the wAIC were significantly smaller (worse) for the bSync model (*M* = .12, *SD* = .026) than for the RW model (*M* = .78, *SD* = .027). A second group (7 subjects) showed wAIC values that were approximately equally strong for the bSync (*M* = .44, *SD* = .036) as for the RW model (*M* = .50, *SD* = .032). In a third group (12 subjects), the bSync model showed wAIC that were significantly higher for the bSync model (*M* = .64, *SD* =. 027) than for the RW model (*M* = .32, *SD* = .026).

**Table 1.**
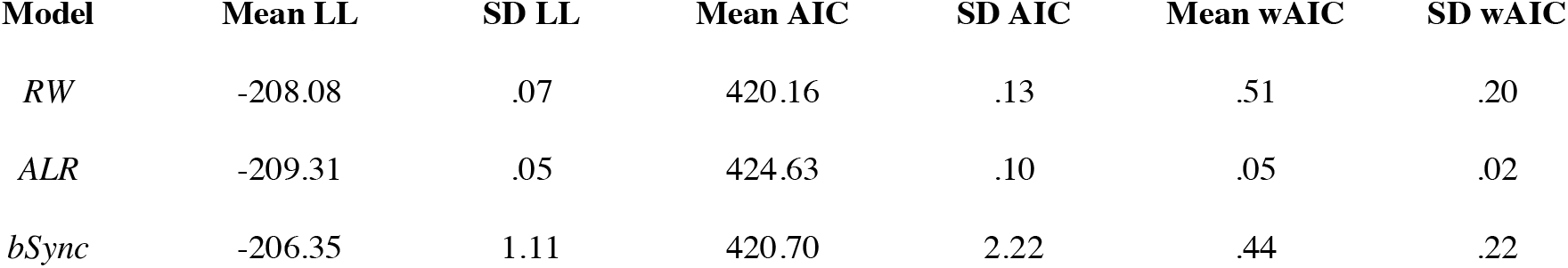
Goodness of fit measures. Results of log-likelihood (LL), AIC and wAIC computations over subjects are shown for each of three models. For LL and wAIC, high values indicate a better fit, while for AIC a low value indicates a good fit.

| Model | Mean LL | SD LL | Mean AIC | SD AIC | Mean wAIC | SD wAIC |
| --- | --- | --- | --- | --- | --- | --- |
| *RW* | -208.08 | .07 | 420.16 | .13 | .51 | .20 |
| *ALR* | -209.31 | .05 | 424.63 | .10 | .05 | .02 |
| *bSync* | -206.35 | 1.11 | 420.70 | 2.22 | .44 | .22 |

**Fig 2.**
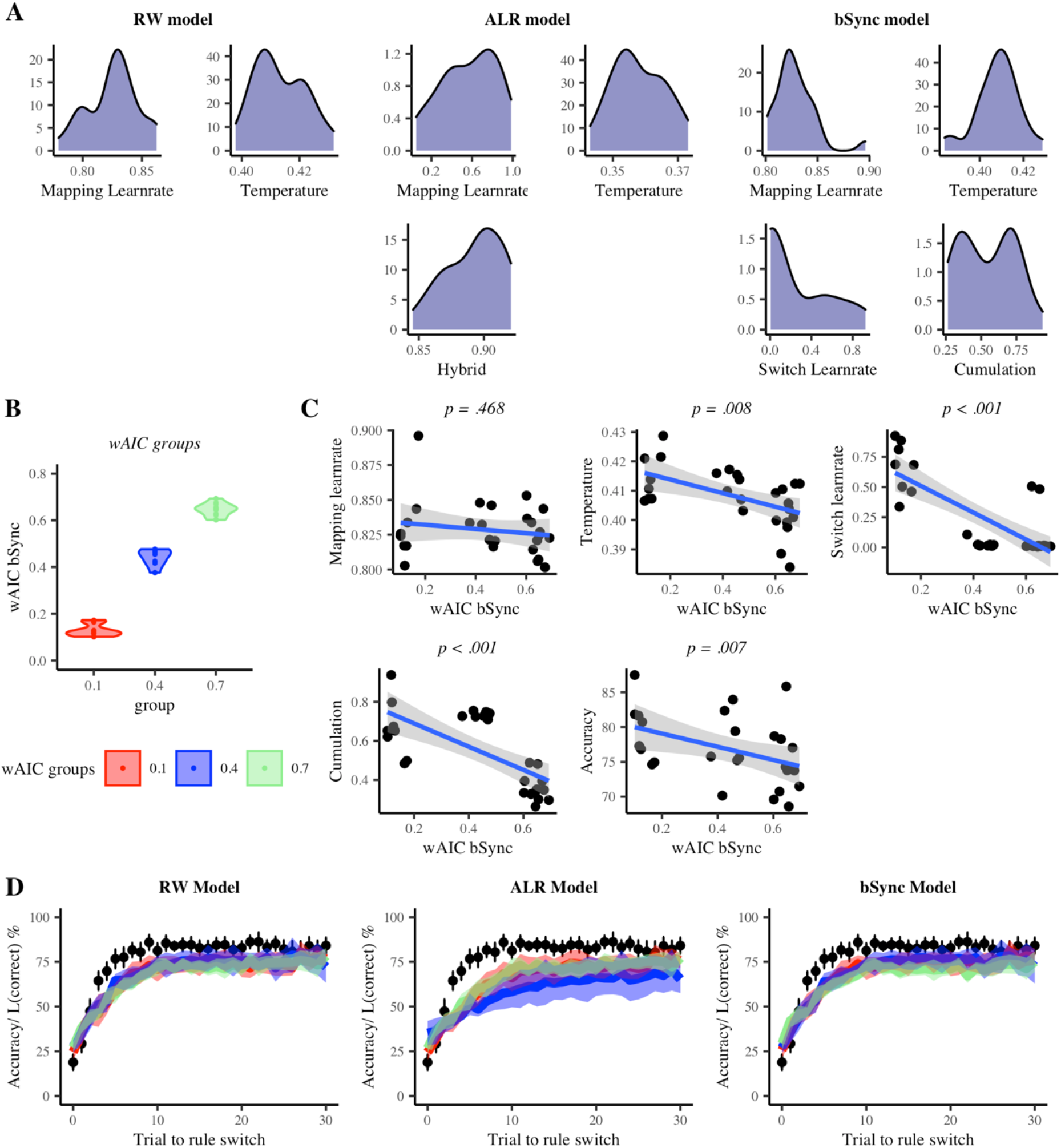
Model comparison. *A: Parameter distributions*. Distributions of fitted parameter values are shown for each model. *B: wAIC groups*. This figure illustrates how wAIC values can be roughly divided in three groups (colors). *C: Correlation plots*. Correlations are shown between wAIC of the bSync model and all parameters of the bSync model. In the lower middle plot, also the correlation between wAIC and task accuracy is shown. *D: Learning curve fit*. Black dots represent the mean accuracy data over all subjects. The error bars show the 95% confidence intervals. The colored lines illustrate the mean Likelihood of the correct response for each wAIC group in B. The shades represent the 95% confidence interval.

Three parameters of the bSync model showed a significant correlation with wAIC (Fig 2C). These parameters were the Switch learning rate (*rho* = −.761, *p* < .001), the Cumulation parameter (*rho* = −.708, *p* < .001), and the Temperature parameter (*rho* = −.497, *p* = .008). There was no significant correlation with the Mapping learning rate (*rho* = −.145, *p* = .468). Additionally, a correlation test between accuracy and wAIC revealed that the bSync model fitted significantly better for subjects with a lower accuracy (*rho* = −.510, *p* = .007). Also correlations between wAIC values and parameters of the other two models were tested but none of these correlations reached significance.

We next estimated a learning curve for each model and each wAIC group (Fig 2D). This learning curve represents the estimated likelihood of the correct response averaged over all rule switches and all subjects within a group. Differences in learning curve between the three groups are very subtle: We conclude that an average measure like switch-locked learning curve does not suffice to empirically distinguish between the three models.

As described in Equation (5), the bSync model only uses negative prediction errors to evaluate rule switches. Nevertheless, it might be argued that also positive prediction errors determine rule switching. To test this, an alternative version of the bSync model (bSync-linear) was currently also fitted. Here, f(*Rew* − *V*(*R*)) = −(*Rew* − *V*(*R*)) for all trials. Hence, switch evidence increased for negative prediction errors and decreased for positive prediction errors. Here, we observed a clear advantage in terms of AIC for the original bSync model (*M* = 420.70, *SD* = 2.22) compared to the alternative bSync-linear model (*M* = 481.12, *SD* = 75.28). Hence, only the original bSync model was used for the consecutive analyses.

In sum, we found that, for the bSync model, participants’ behavior is best explained by the model version that is biased towards negative prediction errors to evaluate rule switches. When comparing this bSync model with the RW and ALR models, three groups of participants could be distinguished. Moreover, the individual measures of model fit correlated significantly with accuracy and several parameters of the bSync model.

### EEG and Model Data

#### Power Cluster Analyses

Cluster analysis on post-feedback power revealed three significant clusters that were selective for feedback processing (Fig 3). All three clusters appeared between 0 and 750 ms from feedback onset. As was predicted by the Sync model (Fig 3A), one of these clusters was in the theta frequency range (∼ 4-8 Hz) and located on midfrontal electrodes (Fig 3B, D). This theta cluster showed more power for negative than for positive feedback. Additionally, we found two clusters located on the posterior channels. One of these clusters was in the delta frequency (< 4 Hz; Fig 3B, E), the other cluster was located in the alpha-frequency range (∼ 8-15 Hz; Fig 3B, C). Both the delta and alpha cluster showed less power for negative feedback than for positive feedback. No correlation between the power contrast of a cluster and subjects’ wAIC for the bSync reached significance.

**Fig 3.**
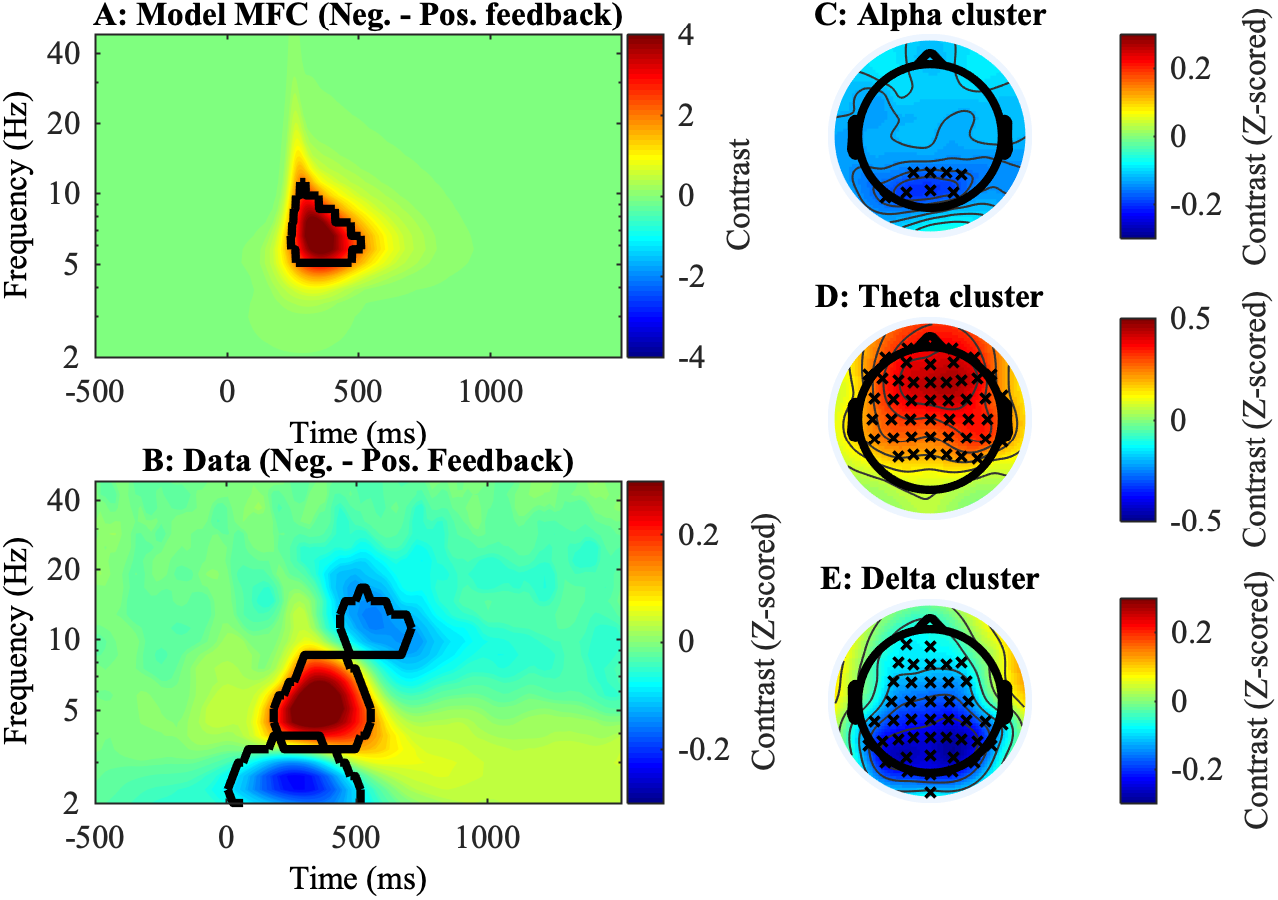
Power results. *A-B: Time-Frequency plots of contrast (Negative – Positive feedback)*. Significant clusters are indicated by the black contour line. *A:* Contrast of power in the model pMFC. *B:* Contrast for Z-scored power in the human data, averaged over all 64 electrodes. *C-E: topographical plots of clusters found in the human data*. Crosses indicate channels where the contrast reached significance.

#### Midfrontal Theta Power and Prediction Error

We next consider the first of three model-driven EEG hypotheses. We first perform statistical analysis on the Sync-model simulated data (Fig 4A). Theta power in the Sync model data was best predicted by the regression model that included an interaction between reward and prediction error (*F*(1, 11980) = 22133, *p* < .001). Hence, there was a significant main effect of prediction error (*F*(1, 11980) = 742962, *p* < .001, *β* = −4.99) and a significant interaction of prediction error and reward (*F*(1, 11980) = 22133, *p* < .001, *β* = 4.48). Thus, as predicted, the model cluster showed a negative linear relationship with negative prediction error, and no linear relationship with positive prediction error (Fig 4A).

**Fig 4.**
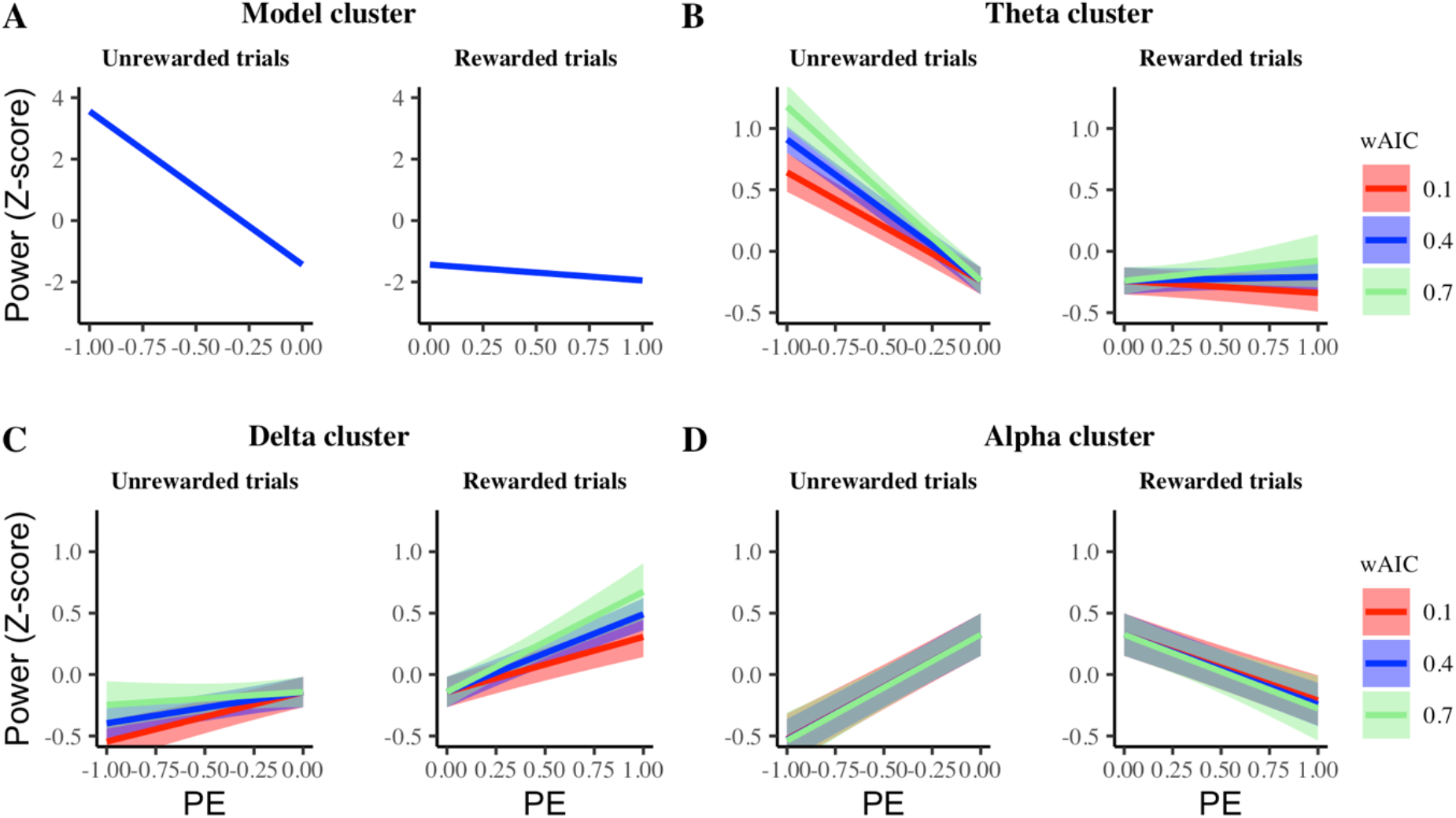
Result of linear regression between power and prediction error (PE) in all clusters. Lines illustrate the trial-by-trial relation between the estimated prediction errors and the mean power extracted from the clusters in Fig 3. The shades represent 95% confidence intervals. The Model cluster (A) aimed to predict empirical data from the theta cluster (B). For exploratory purposes, also the relation between estimated prediction errors and power in the delta (C) and alpha (D) cluster are shown.

In order to test this prediction in the empirical theta cluster (cluster reported in the previous section), prediction errors were estimated by simulating the bSync model (see Fig 5A). Importantly, these prediction error estimates were extracted from learning in the Switch unit (see Equation (4)) and not from the learning of stimulus-action pairs in the Mapping unit (Equation (3). For theta power, the regression model including the interaction between prediction error and reward fitted significantly better than the regression model with only prediction error as regressor (*χ*^2^ (1, *N* = 27) = 110, *p* < .001). Additionally, the regression model including the three-way interaction between prediction error, reward and wAIC fitted significantly better than the regression model with only the two-way interaction (*χ*^2^ (2, *N* = 27) = 20.74, *p* < .001). Here, all effects reached significance. Hence, there was a main effect of prediction error (*χ*^2^ (1, *N* = 27) = 1299, *p* < .001, *β* = −.79) and an interaction of prediction error with reward (*χ*^2^ (1, *N* = 27) = 110, *p* < .001, *β* = .65). Additionally, there was a significant interaction between prediction error, reward and wAIC (*χ*^2^ (2, *N* = 27) = 20.90, *p* < .001). As can be observed in Fig 4B these results indicated a significant negative linear relationship between power and negative prediction error, which was stronger for subjects with a high wAIC (i.e., better behavioral fit of the Sync model); and an absence of linear relationship between power and positive prediction error which did not differ significantly for wAIC (Fig 4B). Interestingly, the three-way interaction was significant in the unrewarded (negative prediction error) trials (*β* = −.89, *p* < .001) but did not reach significance in the rewarded (positive prediction error) trials (*β* = .44, *p* = .077).

**Fig 5.**
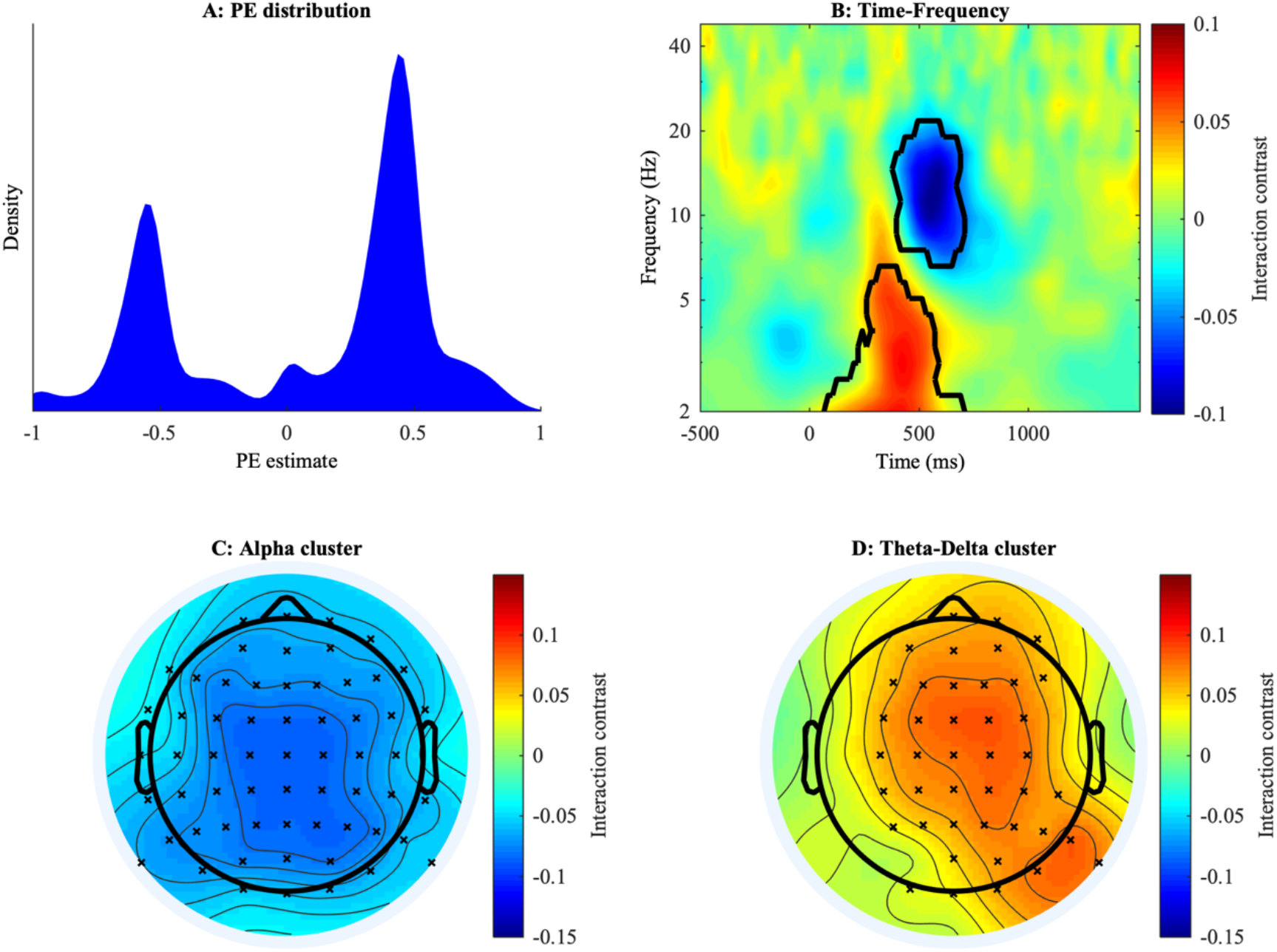
Interaction between prediction error and reward in power. *A: Distribution of prediction error estimates*. Note that these prediction error estimates are not used for learning stimulus-action pairs, but for module learning in the Switch unit (Equation (4)). *B: Time-frequency results*. Colors represent the contrast value of the interaction effect. Black contours indicate significant clusters. *C: Topography of the alpha interaction cluster. D: Topography of the theta-delta interaction cluster*. Crosses indicate channels where the contrast reached significance.

For exploratory purposes, we investigated the same regression models in the delta and alpha clusters. In the delta cluster, the difference in regression model fit between the regression models without and with the prediction error-reward interaction term did not reach significance (*χ*^2^ (1, *N* = 27) = 3.49, *p* = .062). However, the regression model that also included the three-way interaction between prediction error, reward and wAIC fitted significantly better than the regression model with no interaction terms (*χ*^2^ (3, *N* = 27) = 9.27, *p* = .026). Here, the main effect of prediction error was significant (*χ*^2^ (1, *N* = 27) = 580, *p* < .001, *β* = .45). The interaction between prediction error and reward did not reach significance (*χ*^2^ (1, *N* = 27) = 3.49, *p* = .062, *β* = −.07). Also the three-way interaction term did not reach significance (*χ*^2^ (2, *N* = 27) = 5.83, *p* = .054). However, if the interaction was considered separately for rewarded trials (*β* = .61, *p* = .018) and unrewarded trials (*β* = −.50, *p* = .033), both reached significance. As can be observed in Fig 4C, this meant that there was a positive linear relationship between power and prediction error for both positive and negative prediction error (Fig 4C). For subjects with low wAIC, the slope in unrewarded trials was similar to the slope in rewarded trials, while for subjects with high wAIC, an inverse effect of the theta cluster was observed in which there was a flat slope in unrewarded trials but a steeper slope in rewarded trials.

In the alpha cluster, the regression model with the two-way interaction term showed a significantly better fit than the regression model without interaction (*χ*^2^ (1, *N* = 27) = 224, *p* < .001). When the three-way interaction was added, it did not lead to a significantly better regression model (*χ*^2^ (2, *N* = 27) = .35, *p* = .841). Here, a significant main effect of prediction error (*χ*^2^ (1, *N* = 27) = 142, *p* < .001, *β* = .85) and a significant interaction between prediction error and reward (*χ*^2^ (1, *N* = 27) = 226, *p* < .001, *β* = −1.38) were observed. The three-way interaction between prediction error, reward and wAIC was not significant (*χ*^2^ (2, *N* = 27) = .360, *p* = .833). As is shown in Fig 4D, power in the alpha cluster exhibited a positive linear relationship for negative prediction error, but a negative linear relationship with positive prediction error. These effects did not differ with respect to wAIC.

To explore the topology of these interaction effects described above, we conducted another cluster analysis. Here, we multiplied prediction error (scaled separately for positive and negative prediction errors) with reward (−1 for unrewarded trials and 1 for rewarded trials) as a regressor for power. This resulted in a contrast value for the interaction between prediction error and reward for each electrode, timepoint and frequency. These contrast values were then entered into the clustering algorithm. As expected, we observed a significant cluster in the alpha frequency (Fig 5B) which was strongest on posterior electrodes (Fig 5C). We also observed significant effects in the theta and delta frequency ranges (Fig 5D). Although the interaction pattern for theta (Fig 4B) and delta (Fig 4C) are mirrored (and thus qualitatively different), they are represented by a similar contrast value, because in both theta and delta empirical patterns, the slope for positive prediction errors is larger than the slope for negative prediction errors. Because they are also topographically (partially) overlapping, they were clustered together by the algorithm, resulting in one cluster that was a mixture of the theta and delta effects on both time-frequency and topographical level.

In sum, in line with model predictions, we found a linear relationship between post-feedback theta power and negative prediction errors but not with positive prediction errors (interaction between reward and prediction error). Moreover, we found that this interaction effect was stronger for participants that fitted better with the bSync model. On top of model predictions, two other clusters could be distinguished in post-feedback power. Here, a delta cluster showed an almost exactly mirrored pattern relative to the theta cluster. An alpha cluster showed an inversed U-shaped pattern with respect to prediction errors.

#### Rule Switch Locking

For the second model-driven EEG hypothesis, power from the theta, alpha, and delta clusters was extracted in trials within a 31-trial window from the rule switch (−15 to +15). In all clusters, one trial significantly deviated from baseline power. In the theta cluster (Fig 6A), only the rule switch (0; i.e., all trials exactly at rule switch) was significant above baseline (CI99.84 [−2.059, .256], *baseline* = −2.340). Linear regression of the data time course (across 31 trials) on the Sync model time course showed a significant effect (*F*(1, 835) = 20.51, *p* < .001, *R*^2^ _adj_ = .023, *β* = .31). In the delta cluster (Fig 6B), only the rule switch (0) was significantly below baseline (CI99.84 [−2.450, −1.265], *baseline* = −1.201). Linear regression of the data time course on the Sync model time course revealed a significant correlation (*F*(1, 835) = 7.36, *p* = .007, *R*^2^ _adj_= .008, *β* = −.18). For the alpha cluster (Fig 6C), again one trial was significantly below baseline (CI99.84 [−6.275, −3.603], *baseline* = −3.584). Notably, this was the point right after the rule switch (+1; i.e., all trials right after the rule switch). Moreover, when data was locked to the moment where subjects indicated the rule switch (Fig 6D), alpha power reaches a minimum at this exact moment (CI99.84 [−7.675, −3.686], *baseline* = −3.584). Also in the alpha cluster, the linear regression of the power on the Sync model pattern reached significance with a negative slope (*F*(1, 835) = 32.72, *p* < .001, *R*^2^ _adj_ = .037, *β* = −.65).

**Fig 6.**
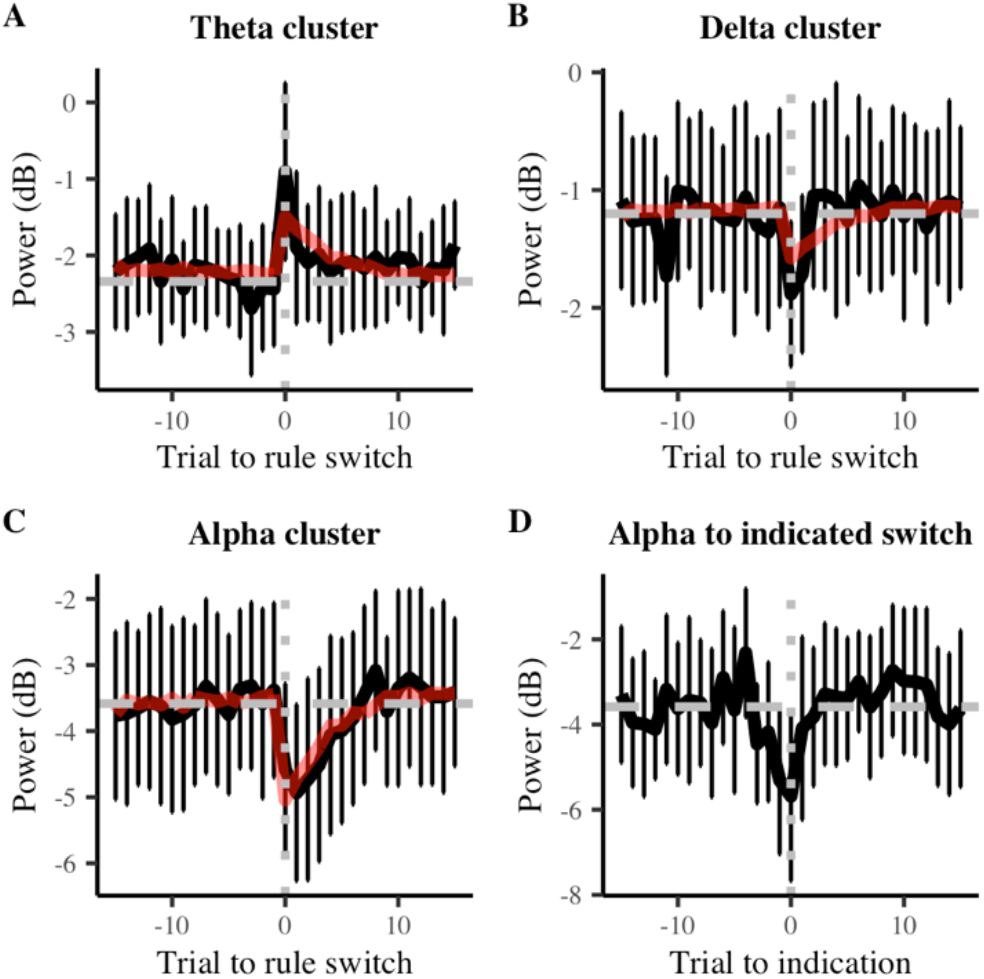
Power locked to rule switch. Black lines show the mean power. Error bars show the 99.84% confidence interval (Bonferroni correction). The horizontal grey dashed line represents baseline power and the vertical grey dotted line indicates the moment of the rule switch. The red line visualizes the result of linear regression between the Sync model and human data. *A-C* show data locked to the moment of the actual rule switch. *D* shows data of the alpha cluster locked to the moment when subjects indicated they noticed the task switch.

Power at the peak trials (trials at point 0 for theta and delta, trials at point +1 for alpha) was extracted and added to a linear regression with wAIC as predictor. This revealed no significant effects for the theta (*F*(1, 25) = .004, *p* = .948, *R*^2^ _adj_= −.040, *β* = −.10) or delta cluster (*F*(1, 25) = .680, *p* = .417, *R*^2^ _adj_ = −.012, *β* = .66). However, the effect of wAIC did reach significance in the alpha cluster (*F*(1, 25) = 7.22, *p* = .013, *R*^2^ _adj_= .193, *β* = 4.17). Fig 7 sheds light on how activity in the alpha cluster differed depending on wAIC. For illustrative purposes, subjects were divided in three groups of low, middle and high wAIC. For each group, the data pattern of alpha activity was plotted, once locked to the real rule switch (Fig 7A) and once locked to the indication of a rule switch (Fig 7B). Here, it is observed that the alpha pattern is mainly driven by subjects with a low wAIC (i.e., bad fit) for the bSync model.

**Fig 7.**
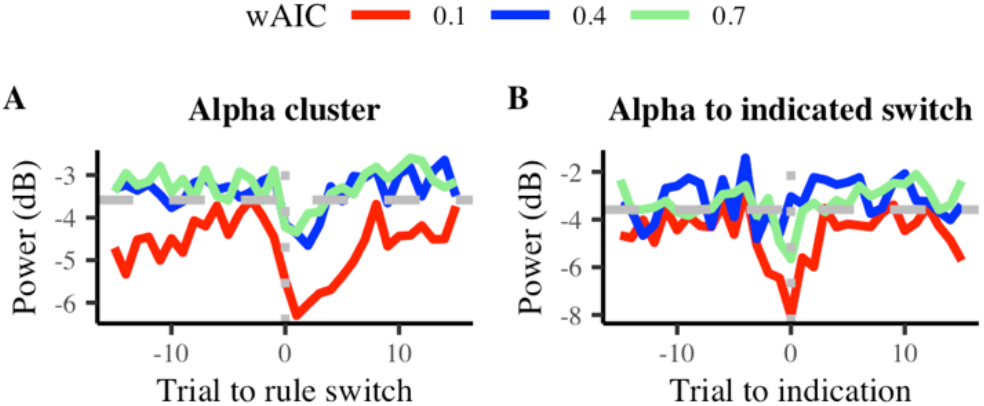
Power locked to rule switch for different wAIC. Data patterns are shown for different wAIC values (colored lines). The horizontal grey dashed line shows the baseline power over all subjects and the vertical grey dotted line indicates the moment of the rule switch (A) or indication of rule switch (B).

In sum, simulated theta power significantly predicted empirical theta power. Here, theta power peaked at the moment of a rule switch. Just like in the first model-driven EEG hypothesis, power from the empirical delta cluster showed the mirrored pattern compared to theta. Remarkably, alpha cluster showed a dip in power, not with respect to the actual rule switch but with respect to the subjectively indicated rule switch.

#### Midfrontal-Posterior Phase-Coupling Analyses

We next turn to our third model-driven EEG analysis concerning an increase of phase-coupling between midfrontal and posterior electrodes after negative feedback. As previously described, in the Sync model, this coupling is induced by bursts that are sent from pMFC to posterior areas in the Mapping unit. Since pMFC power is stronger after negative feedback, also the number of bursts and the amount of phase-coupling is increased. To investigate this, we looked at phase-coupling between a midfrontal electrode (FCz) and all lateral electrodes.

Here, non-parametric cluster analyses on the phase-locking data (Fig 8) revealed six significant clusters that were selective for feedback (for details see Materials and Methods). These clusters were located in the theta (4; Fig 8A, B, C) or delta (2; Fig 8A, B, D) frequency band. In the theta frequency band, two clusters were located at temporal electrodes; two other clusters were located on more lateral/anterior frontal electrodes. In the delta frequency band, both clusters were located on posterior electrodes. In line with the results of Sync model simulations (Fig 8E), the theta clusters showed an increase in phase-locking after negative feedback. This was the case for both the ipsilateral and contralateral electrodes. The delta clusters show the inverse pattern of the theta cluster. Here, phase-locking was stronger after positive feedback than after negative feedback in both the ipsi- and contralateral cluster. As in the power analyses, we also explored whether the phase-locking contrast in each cluster correlated with the subjects’ wAIC for the bSync model. None of these correlations reached significance.

**Fig 8.**
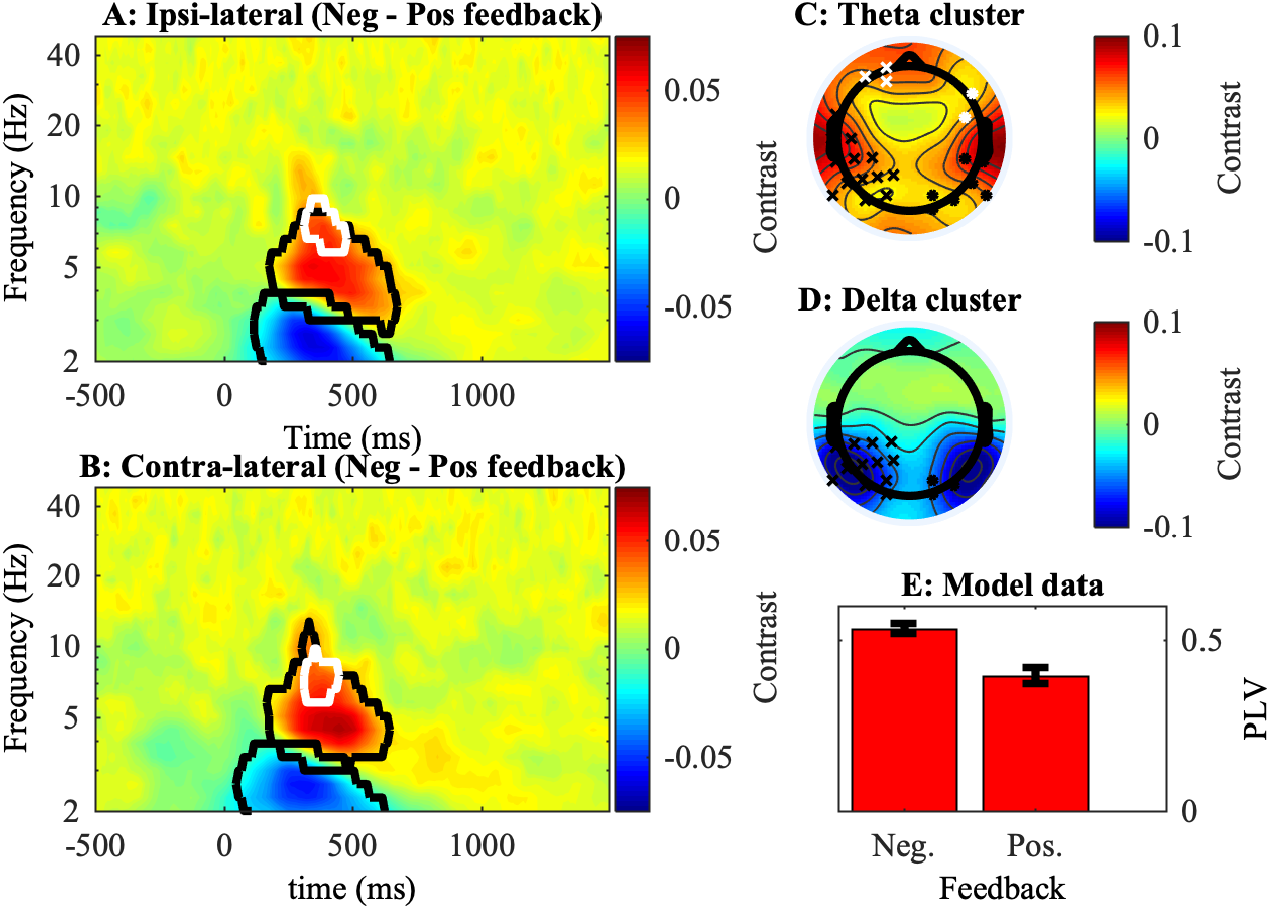
Phase-locking with respect to FCz. *A-B: Time-Frequency plots of contrast (Negative – Positive feedback)*. Significant clusters are indicated by the black or white contour line. The black line represents posterior clusters in C and D (black crosses) while the white line represents the frontal clusters in C (white crosses). All plots show phase-locking with respect to the FCz electrode. A: Contrast of iPLV averaged over all ipsi-lateral electrodes. B: Contrast of iPLV averaged over all contra-lateral electrodes. *C-D: Topographical plots of clusters*. Data was averaged over all time points and frequencies that were included in the respective contours of A and B. Channels where the contrast reached significance are marked by crosses or dots. The left channels (crosses) present ipsi-lateral electrodes and the right channels (dots) present contra-lateral electrodes. Again, the white color was used to distinguish the frontal clusters from the temporal clusters. *E:* Predicted phase-coupling in the model for the 250-500 ms post-feedback period.

In sum, as predicted by the Sync model, we found stronger phase-coupling in the theta frequency between midfrontal and more posterior electrodes after negative feedback than after positive feedback. Additionally, we found an increase in phase-coupling between midfrontal and lateral frontal electrodes, also in the theta frequency range. Similar to our power analyses, we found an inverse effect in the delta frequency compared to the theta frequency.

## Discussion

The current study aimed to gain insight in the neural mechanisms that allow humans to flexibly adapt to rule changes in the environment. Twenty-seven healthy human subjects were tested on a probabilistic reversal learning task while measuring EEG. Behaviorally, three models of increasing hierarchical complexity were compared. A first, RW model, updated the value of stimulus-action mappings on a trial-by-trial basis with a fixed learning rate. In a second, ALR model, this approach was extended with an adaptable learning rate, allowing the ALR model to flexibly adapt to rule switches (fast learning) but to also be robust to noise evoked by probabilistic feedback (slow learning). The third, Sync model implemented modularity to retain task-specific mappings. It employs hierarchical learning to determine when to switch between rule modules. No evidence was found for the ALR model, while for some subjects the RW model fit best, and for others the Sync model.

Simulations of the Sync model allowed formulation and testing of three model-driven EEG hypotheses. The first hypothesis concerns midfrontal theta and prediction errors. In the Sync model, prediction errors are used to evaluate how much control is needed. The level of control is represented by theta power in the pMFC. Since only negative prediction errors inform about rule switches, the Sync model increased control after negative prediction errors but not after positive prediction errors. Thus, although most previous work (Cavanagh, Cohen, & Allen, 2009; Cavanagh, Frank, Klein, & Allen, 2010; Ergo, De Loof, Janssens, & Verguts, 2019) described a U-shape relationship between prediction error and theta power, we currently hypothesized and observed a selectivity for negative prediction errors (see also Janssen et al., 2016). A linear relationship between prediction error and power in the theta cluster was observed for unrewarded trials (negative prediction error) but not for rewarded trials (positive prediction error). This effect was stronger for subjects with a better Sync model fit. Based on our theoretically driven hypothesis, we did not extract prediction errors from stimulus-action learning but from learning in the Switch unit. Future research should investigate how midfrontal theta is influenced by different types of prediction errors.

Since prediction errors are strongest at the rule switch, a second model-driven hypothesis stated that theta power peaks at rule switches. Again, this hypothesis was empirically supported. Moreover, simulated power significantly predicted power in the empirical theta cluster. Consistent with earlier work (Cunillera et al., 2012; Sauseng et al., 2006), theta power increased and alpha power decreased at rule switches. How this theta increase relates to the alpha decrease, and to the individual differences that we observed, deserves future research.

Current work provides a mechanistic explanation how increases in theta power after prediction errors implement new task rules by synchronizing modules. This resulted in a third model-driven hypothesis. Here, the Sync model uniquely predicted that phase connectivity would increase after negative feedback. We found six significant clusters. Four of them were in theta frequency range and showed the predicted pattern. Two of these clusters were located on posterior-temporal electrodes, roughly in line with our prediction of motor and visual areas. The remaining four clusters were consistent with previous work (Cavanagh et al., 2010) showing a feedback-locked, prediction-error induced increase of theta phase-coupling between midfrontal and lateral frontal sites, and a delta coupling decrease between midfrontal and posterior cortical sites.

Several hypotheses remain to be tested. First, as mentioned in the Methods, previous modeling work (Verbeke & Verguts, 2019) used gamma frequency in the Mapping unit instead of theta frequency. This frequency was currently changed because empirical work demonstrated within-frequency (theta-theta) coupling (Cavanagh et al., 2009; Clouter, Shapiro, & Hanslmayr, 2017) during cognitive tasks, in addition to cross-frequency coupling. We thus also studied within-frequency coupling empirically. Nevertheless, future work, using MEG or more invasive measurements, should also study the role of cross-frequency (theta-gamma) coupling. Second, the limited spatial resolution of EEG did not allow testing whether task rules are implemented by synchronizing task-relevant modules.

Several model extensions can be made. For instance, while for the current reversal learning task it was sufficient to use prediction error to determine when to make a binary switch, a more sophisticated approach might apply in everyday life, where contextual cues allow navigating a vast map of tasks and rules. One way to address this issue is by adding second-level contextual features which allow the LFC to (learn to) infer which of multiple task modules should be synchronized. Additionally, scalability of the Sync model is currently limited by how modularity was implemented in the Mapping unit. Here, none of the rule-1 mappings are shared with rule 2. Such a strict division of task mappings is optimal when mappings are orthogonal. However, when some mappings can be generalized between tasks, the current approach does not allow knowledge transfer across contexts. As addressed earlier (Collins & Frank, 2013; Gershman, Blei, & Niv, 2010), a more sustainable way is to construct modules of mappings that are shared between tasks. Instead of learning each new task from scratch, this approach allows transferring partial knowledge between tasks. Future work should explore whether these more complex hierarchical learning algorithms can be integrated in the Sync model.

Recent work emphasized that reinforcement learning can operate not only over observed states, but also over belief states that an agent may infer (Gershman & Uchida, 2019; Wilson et al., 2014). In the Sync model, there were no (contextual) cues. Therefore, the Sync model could rely exclusively on prediction errors to estimate the belief state (task rule) of the environment. When contextual features are added, a future version of the Sync model may estimate belief states in a more efficient manner. Note also that the Sync model uses two types of prediction error: One to adjust lower-level mappings, and another to determine the (higher-level) task rule state. Instead, non-hierarchical models (e.g., RW, ALR) use prediction errors only to adjust lower-level mappings.

Building on suggestions of previous work (Piray, Dezfouli, Heskes, Frank, & Daw, 2019), the current study illustrated how individual differences in model fit can be leveraged to address cognitive questions. Three groups were distinguished: one group aligned with the RW model, a second group aligned with the Sync model and in a third group, the RW and Sync model could not be empirically distinguished. Although the differences between groups were non-significant when averaging over several trials (e.g., learning curve), more fine-grained measures (e.g., wAIC, trial-by-trial power) revealed important individual differences. Interestingly, subjects with lower accuracy fitted better with the Sync model. This is consistent with previous work (Verbeke & Verguts, 2019) which illustrated that modularity as employed by the Sync model is only beneficial if the learning problem is sufficiently complex. Furthermore, despite previous work showing a good behavioral fit of the ALR model (Bai et al., 2014), the fit of the ALR model in the current study was consistently low. In contrast to previous studies, the current task applied more frequent task rule switches without long stable trial blocks, favoring constant high learning rates. Thus, future work should investigate whether subjects employ the RW, ALR, or Sync framework depending on the structure and complexity of the task.

The Sync model implements modularity via neural oscillations between task-relevant areas. This concords with a role of neural oscillations for a wide variety of cognitive functions, including visual attention (Gray & Singer, 1989; Jensen, Bonnefond, & VanRullen, 2012), working memory (Hsieh, Ekstrom, & Ranganath, 2011; Lisman & Idiart, 1995), cognitive control (Cavanagh & Frank, 2014) and declarative learning (Ergo, De Loof, & Verguts, 2020). According to the BBS hypothesis (Fries, 2015), these cognitive functions require binding of several stimuli or features. Current work described how oscillations, and more specifically synchronization, might be relevant in hierarchical rule learning.

On anatomical-functional level, we built on suggestions from previous work that pMFC cooperates with LFC to exert hierarchical control over lower-level motor processes (Alexander & Brown, 2015; Koechlin, Ody, & Kouneiher, 2003). In the Sync model, LFC signals which rule modules should be synchronized. Consistently, previous theories describe LFC as containing task demands (Botvinick et al., 2001), and empirical work found strong communication between LFC and pMFC in cognitive tasks (Cavanagh et al., 2010; Kondo, Osaka, & Osaka, 2004; Mac Donald, Cohen, Stenger, & Carter, 2000). Also in line with previous data (Boorman, Behrens, Woolrich, & Rushworth, 2009; Holroyd & McClure, 2015; Wilson et al., 2014), the model aMFC keeps track of the relevant task rule. Additionally, consistent with fMRI work (Aben, Calderon, Van den Bussche, & Verguts, 2020), current study found increased coupling between midfrontal cortex and task-related areas when more control was needed (negative feedback). While this fMRI work showed anatomically detailed networks of connectivity, current study described how this connectivity may work at algorithmic level.

To summarize, we have demonstrated how the brain might employ synchronization to bind task-relevant areas for efficient rule switching. To achieve this, we used EEG, computational modelling, individual differences, and behavioral analysis. We believe that this approach might reveal how more complicated tasks can be implemented via synchronization as well.

## Acknowledgements

PV and KE were supported by grant 1102519N and grant 1153418N from Research Foundation Flanders, respectively. EDL and TV were supported by grant BOF17/GOA/004 from the Ghent University Research Council. We thank Clay Holroyd for useful comments on this paper.

## References

Aben, B., Calderon, C. B., Van den Bussche, E., & Verguts, T. (2020). Cognitive effort modulates connectivity between dorsal anterior cingulate cortex and task-relevant cortical areas. The Journal of Neuroscience, 40(19), JN-RM-2948-19. https://doi.org/10.1523/jneurosci.2948-19.2020

Alexander, W. H., & Brown, J. W. (2015). Hierarchical error representation: A computational model of anterior cingulate and dorsolateral prefrontal cortex. Neural Computation, 27(11), 2354–2410. https://doi.org/10.1162/NECO

Bai, Y., Katahira, K., & Ohira, H. (2014). Dual learning processes underlying human decision-making in reversal learning tasks: Functional significance and evidence from the model fit to human behavior. Frontiers in Psychology, 5(AUG), 1–8. https://doi.org/10.3389/fpsyg.2014.00871

Behrens, T. E. J., Woolrich, M. W., Walton, M. E., & Rushworth, M. F. S. (2007). Learning the value of information in an uncertain world. Nature Neuroscience, 10(9), 1214–1221. https://doi.org/10.1038/nn1954

Boorman, E. D., Behrens, T. E. J., Woolrich, M. W., & Rushworth, M. F. S. (2009). How green is the grass on the other side? Frontopolar cortex and the evidence in favor of alternative courses of action. Neuron, 62(5), 733–743. https://doi.org/10.1016/j.neuron.2009.05.014

Botvinick, M. M., Braver, T. S., Barch, D. M., Carter, C. S., & Cohen, J. D. (2001). Conflict monitoring and cognitive control. Psychological Review, 108(3), 624–652. https://doi.org/10.1037/0033-295X.108.3.624

Bruña, R., Maestú, F., & Pereda, E. (2018). Phase locking value revisited: Teaching new tricks to an old dog. Journal of Neural Engineering, 15(5). https://doi.org/10.1088/1741-2552/aacfe4

Cavanagh, J. F., Cohen, M. X., & Allen, J. J. B. B. (2009). Prelude to and resolution of an error : EEG phase synchrony reveals cognitive control dynamics during action monitoring. Journal of Neuroscience, 29(1), 98–105. https://doi.org/10.1523/JNEUROSCI.4137-08.2009

Cavanagh, J. F., & Frank, M. J. (2014). Frontal theta as a mechanism for cognitive control. Trends in Cognitive Sciences, 18(8), 414–421. https://doi.org/10.1016/j.tics.2014.04.012

Cavanagh, J. F., Frank, M. J., Klein, T. J., & Allen, J. J. B. B. (2010). Frontal theta links prediction errors to behavioral adaptation in reinforcement learning. NeuroImage, 49(4), 3198–3209. https://doi.org/10.1016/j.neuroimage.2009.11.080

Clouter, A., Shapiro, K. L., & Hanslmayr, S. (2017). Theta phase synchronization is the glue that binds human associative memory. Current Biology, 27(20), 1–6. https://doi.org/10.1016/j.cub.2017.09.001

Cohen, J. D., Dunbar, K., & McClelland, J. L. (1990). On the control of automatic processes: a parallel distributed processing account of the Stroop effect. Psychological Review, 97(3), 332–361. https://doi.org/10.1037/0033-295X.97.3.332

Cohen, M. X. (2014). Analyzing neural time series data: Theory and practice. Cambridge, Massachusetts: The MIT Press.

Collins, A. G. E., & Frank, M. J. (2013). Cognitive control over learning: Creating, clustering, and generalizing task-set structure. Psychological Review, 120(1), 190–229. https://doi.org/10.1037/a0030852

Cools, R., Clark, L., Owen, A. M., & Robbins, T. W. (2002). Defining the neural mechanisms of probabilistic reversal learning using event-related functional magnetic resonance imaging. The Journal of Neuroscience : The Official Journal of the Society for Neuroscience, 22(11), 4563–4567. https://doi.org/20026435

Cunillera, T., Fuentemilla, L., Periañez, J., Marco-Pallarès, J., Krämer, U. M., Càmara, E., … Antoni, R. F. (2012). Brain oscillatory activity associated with task switching and feedback processing. Cognitive, Affective and Behavioral Neuroscience, 12(1), 16–33. https://doi.org/10.3758/s13415-011-0075-5

Delorme, A., & Makeig, S. (2004). EEGLAB: An open source toolbox for analysis of single-trial EEG dynamics including independent component analysis. Journal of Neuroscience Methods, 134(1), 9–21. https://doi.org/10.1016/j.jneumeth.2003.10.009

Ergo, K., De Loof, E., Janssens, C., & Verguts, T. (2019). Oscillatory signatures of reward prediction errors in declarative learning. NeuroImage, 186(September 2018), 137–145. https://doi.org/10.1016/j.neuroimage.2018.10.083

Ergo, K., De Loof, E., & Verguts, T. (2020). Reward prediction error and declarative memory. Trends in Cognitive Sciences, 24(5), 388–397. https://doi.org/10.1016/j.tics.2020.02.009

French, R. M. (1999). Catastrophic forgetting in connectionist networks. Trends in Cognitive Sciences, 6613(April), 128–135.

Fries, P. (2005). A mechanism for cognitive dynamics: neuronal communication through neuronal coherence. Trends in Cognitive Sciences, 9(10), 474–480. https://doi.org/10.1016/j.tics.2005.08.011

Fries, P. (2015). Rhythms for cognition: Communication through coherence. Neuron, 88(1), 220–235. https://doi.org/10.1016/j.neuron.2015.09.034

Gershman, S. J., Blei, D. M., & Niv, Y. (2010). Context, learning, and extinction. Psychological Review, 117(1), 197–209. https://doi.org/10.1037/a0017808

Gershman, S. J., & Uchida, N. (2019). Believing in dopamine. Nature Reviews Neuroscience, 20(11), 703–714. https://doi.org/10.1038/s41583-019-0220-7

Gray, C. M., & Singer, W. (1989). Stimulus-specific neuronal oscillations in orientation columns of cat visual cortex. Proceedings of the National Academy of Sciences of the United States of America, 86(5), 1698–1702. https://doi.org/10.1073/pnas.86.5.1698

Holroyd, C. B. (2016). The waste disposal problem of effortful control. In T. S. Braver (Ed.), Motivation and cognitive control(pp. 235–260). Hove, UK: Psychology Press.

Holroyd, C. B., & McClure, S. M. (2015). Hierarchical control over effortful behavior by rodent medial frontal cortex: A computational model. Psychological Review, 122(1), 54–83. https://doi.org/10.1037/a0038339

Hsieh, L. T., Ekstrom, A. D., & Ranganath, C. (2011). Neural oscillations associated with item and temporal order maintenance in working memory. Journal of Neuroscience, 31(30), 10803–10810. https://doi.org/10.1523/JNEUROSCI.0828-11.2011

Izquierdo, A., Brigman, J. L., Radke, A. K., Rudebeck, P. H., & Holmes, A. (2017). The neural basis of reversal learning: An updated perspective. Neuroscience, 345, 12–26. https://doi.org/10.1016/j.neuroscience.2016.03.021

Janssen, D. J. C., Poljac, E., & Bekkering, H. (2016). Binary sensitivity of theta activity for gain and loss when monitoring parametric prediction errors. Social Cognitive and Affective Neuroscience, 11(8), 1280–1289. https://doi.org/10.1093/scan/nsw033

Jasper, H. (1958). The ten twenty electrode system of the international federation. Electroencephalogr. Clin. Neurophysiol., (10), 371–375.

Jensen, O., Bonnefond, M., & VanRullen, R. (2012). An oscillatory mechanism for prioritizing salient unattended stimuli. Trends in Cognitive Sciences, 16(4), 200–205. https://doi.org/10.1016/j.tics.2012.03.002

Koechlin, E., Ody, C., & Kouneiher, F. (2003). The architecture of cognitive control in the human prefrontal cortex. Science (New York, NY), 302(5648), 1181–1185. https://doi.org/10.1126/science.1088545

Kondo, H., Osaka, N., & Osaka, M. (2004). Cooperation of the anterior cingulate cortex and dorsolateral prefrontal cortex for attention shifting. NeuroImage, 23(2), 670–679. https://doi.org/10.1016/j.neuroimage.2004.06.014

Lachaux, J. P., Rodriguez, E., Martinerie, J., & Varela, F. J. (1999). Measuring phase synchrony in brain signals. Human Brain Mapping, 8(4), 194–208. https://doi.org/10.1002/(SICI)1097-0193(1999)8:4<194::AID-HBM4>3.0.CO;2-C

Lisman, J. E., & Idiart, M. A. P. (1995). Storage of 7 ± 2 short-term memories in oscillatory subcycles. Science, 267(5203), 1512–1515. https://doi.org/10.1126/science.7878473

Mac Donald, A. W., Cohen, J. D., Stenger, A. V., & Carter, C. S. (2000). Dissociating the role of the dorsolateral prefrontal and anterior cingulate cortex in cognitive control. Science, 288(June), 1835–1838. https://doi.org/10.1126/science.288.5472.1835

Makoto, M. (2018). Makoto’s preprocessing pipeline. Retrieved from https://sccn.ucsd.edu/wiki/Makoto%27s_preprocessing_pipeline

Maris, E., & Oostenveld, R. (2007). Nonparametric statistical testing of EEG- and MEG-data. Journal of Neuroscience Methods, 164(1), 177–190. https://doi.org/10.1016/j.jneumeth.2007.03.024

Nolte, G., Bai, O., Wheaton, L., Mari, Z., Vorbach, S., & Hallett, M. (2004). Identifying true brain interaction from EEG data using the imaginary part of coherency. Clinical Neurophysiology, 115(10), 2292–2307. https://doi.org/10.1016/j.clinph.2004.04.029

Peirce, J., Gray, J. R., Simpson, S., MacAskill, M., Höchenberger, R., Sogo, H., … Lindeløv, J. K. (2019). PsychoPy2: Experiments in behavior made easy. Behavior Research Methods, 51(1), 195–203. https://doi.org/10.3758/s13428-018-01193-y

Piray, P., Dezfouli, A., Heskes, T., Frank, M., & Daw, N. (2019). Hierarchical Bayesian inference for concurrent model fitting and comparison for group studies. PLoS Computational Biology, 15(6), 34. https://doi.org/ https://doi.org/10.1371/journal.pcbi.1007043

R Core Team. (2017). R: A language and environment for statistical computing. Vienna, Austria.

Rescorla, R. A., & Wagner, A. R. (1972). A theory of Pavlovian conditioning: Variations in the effectiveness of reinforcement and nonreinforcement. Classical Conditioning II Current Research and Theory, 21(6), 64–99. https://doi.org/10.1101/gr.110528.110

Saez, A., Rigotti, M., Ostojic, S., Fusi, S., & Salzman, C. D. (2015). Abstract context representations in primate amygdala and prefrontal Cortex. Neuron, 87(4), 869–881. https://doi.org/10.1016/j.neuron.2015.07.024

Sauseng, P., Klimesch, W., Freunberger, R., Pecherstorfer, T., Hanslmayr, S., & Doppelmayr, M. (2006). Relevance of EEG alpha and theta oscillations during task switching. Experimental Brain Research, 170(3), 295–301. https://doi.org/10.1007/s00221-005-0211-y

Silvetti, M., Seurinck, R., & Verguts, T. (2011). Value and prediction error in medial frontal cortex: Integrating the single-unit and systems levels of analysis. Frontiers in Human Neuroscience, 5(August), 75. https://doi.org/10.3389/fnhum.2011.00075

Silvetti, M., Vassena, E., Abrahamse, E., & Verguts, T. (2018). Dorsal anterior cingulate-brainstem ensemble as a reinforcement meta-learner. PLoS Computational Biology, 14(8), 1–32. https://doi.org/10.1371/journal.pcbi.1006370

Springer, M., & Paulsson, J. (2006). Harmonies from noise. Nature, 439(January), 27–29. https://doi.org/doi:10.1038/439027a

The MathWorks Inc. (2016). MATLAB R2016B. Natick, Massachussetts, United States.

Verbeke, P., & Verguts, T. (2019). Learning to synchronize: How biological agents can couple neural task modules for dealing with the stability-plasticity dilemma. PLoS Computational Biology, 15(8). https://doi.org/10.1371/journal.pcbi.1006604

Verguts, T. (2017). Binding by random bursts: A computational model of cognitive control. Journal of Cognitive Neuroscience, 29(6), 1103–1118. https://doi.org/10.1162/jocn

Widrow, B., & Hoff, M. M. E. (1960). Adaptive switching circuits. IRE WESCON Convention Record, 4(1), 96–104.

Wilson, R. C., Takahashi, Y. K., Schoenbaum, G., & Niv, Y. (2014). Orbitofrontal cortex as a cognitive map of task space. Neuron, 81(2), 267–279. https://doi.org/10.1016/j.neuron.2013.11.005

Womelsdorf, T., Johnston, K., Vinck, M., & Everling, S. (2010). Theta-activity in anterior cingulate cortex predicts task rules and their adjustments following errors. Proceedings of the National Academy of Sciences, 107(11), 5248–5253. https://doi.org/10.1073/pnas.0906194107

Womelsdorf, T., Schoffelen, J., Oostenveld, R., Singer, W., Desimone, R., Engel, A. K., & Fries, P. (2007). Modulation of neuronal interactions through neuronal synchronization. Science, 316(1609), 1609–1612. https://doi.org/10.1126/science.1139178

Zhou, T., Chen, L., & Aihara, K. (2005). Molecular communication through stochastic synchronization induced by extracellular fluctuations. Physical Review Letters, 178103(October), 2–5. https://doi.org/10.1103/PhysRevLett.95.178103

